# A conserved local structural motif controls the kinetics of PTP1B catalysis

**DOI:** 10.1101/2023.02.28.529746

**Authors:** Christine Y. Yeh, Jesus A. Izaguirre, Jack B. Greisman, Lindsay Willmore, Paul Maragakis, David E. Shaw

**Affiliations:** D. E. Shaw Research, New York, NY 10036, USA; Department of Biochemistry and Molecular Biophysics, Columbia University, New York, NY 10032, USA

## Abstract

Protein tyrosine phosphatase 1B (PTP1B) is a negative regulator of the insulin and leptin signaling pathways, making it a highly attractive target for the treatment of type II diabetes. For PTP1B to perform its enzymatic function, a loop referred to as the “WPD loop” must transition between open (catalytically incompetent) and closed (catalytically competent) conformations, which have both been resolved by X-ray crystallography. Although prior studies have established this transition as the rate-limiting step for catalysis, the transition mechanism for PTP1B and other PTPs has been unclear. Here we present an atomically detailed model of WPD-loop transitions in PTP1B based on unbiased, long-timescale molecular dynamics simulations and weighted ensemble simulations. We found that a specific WPD-loop region— the PDFG motif—acted as the key conformational switch, with structural changes to the motif being necessary and sufficient for transitions between long-lived open and closed states of the loop. Simulations starting from the closed state repeatedly visited open states of the loop that quickly closed again unless the infrequent conformational switching of the motif stabilized the open state. The functional role of the PDFG motif is supported by the fact that it (or the similar PDHG motif) is conserved across all PTPs. Bioinformatic analysis shows that the PDFG motif is also conserved, and adopts two distinct conformations, in deiminases, and the related DFG motif is known to function as a conformational switch in many kinases, suggesting that PDFG-like motifs may control transitions between structurally distinct, long-lived conformational states in multiple protein families.

## Introduction

Protein tyrosine phosphatase 1B (PTP1B) plays an essential regulatory role in multiple cellular processes, particularly in leptin, insulin, and epidermal growth factor (EGF) signaling pathways—rendering it a highly attractive therapeutic target for diabetes and various forms of cancer.^1^ The catalytic region of PTP1B is highly conserved among all members of the human PTP family^2^ and comprises the PTP loop (containing the nucleophilic C215), the WPD loop (containing the D181 general acid), and the substrate-binding loop (SBL). As with all PTPs, PTP1B catalyzes the hydrolysis of a phosphotyrosine substrate through a phospho-cysteinyl intermediate.^3^ Substrate binding initiates WPD-loop “closure,” leaving D181 poised to activate a water molecule that cleaves the phospho-cysteinyl intermediate^4^ to complete the catalytic cycle.

PTP1B catalysis, like that of other enzymes, requires both fast and slow motions to perform its catalytic function.^5^ The slow transition between the “open” (catalytically incompetent) and “closed” (catalytically competent) conformations of the WPD loop is the rate-limiting step in the catalytic function of both PTP1B and a homologous bacterial protein, YopH (*k_cat_* = 15–60 s^-1^ and *k_cat_* = 700–1000 s^-1^ respectively).^6,7^ Since this discovery, many groups have reported residues, interactions, and secondary structure motifs that are associated with the regulation of PTP1B catalytic activity.^8–12^ Despite such extensive examination of PTP1B, there exists no mechanistic model that describes the structural changes that control the timescale of opening and closing of the WPD loop, which modulates PTP1B’s catalytic activity.

Here we present a computational model of the transition of the WPD loop at an atomic level of resolution; the model reproduces the kinetics of PTP1B and indicates that a short PDFG motif on the WPD loop is responsible for the slow transition. To develop this model, we first used long-timescale molecular dynamics (MD) simulations to sample the slow motion of the PTP1B WPD loop, observing several transitions from the closed to open state. We then used accelerated weighted ensemble^13^ (AWE) simulations to sample transitions both to and from the open state. Our simulations recapitulated the open and closed WPD states and the millisecond-timescale kinetics of the transitions between them. These simulations also produced a set of putative transition state structures, which we further validated and characterized using committor analysis.^14,15^ Finally, using machine learning and feature analysis,^16^ we developed an atomically detailed model of the mechanism of WPD transitions. We found that the PDFG motif acted as a conformational switch, with structural changes in the motif being both necessary and sufficient^17,18^ to distinguish between long-lived open and closed states of the WPD loop, and thus between the states of the PTP1B catalytic cycle.

We then performed a bioinformatic analysis to assess how widely this role of the PDFG motif might be shared. The PDFG motif (or the similar PDHG motif) is known to be conserved in all PTPs^2,19^ (the histidine residue is similarly bulky and aromatic to the Phe182 in PTP1B): This strongly suggests that the function of the PDFG motif is conserved in tyrosine phosphatases. Furthermore, we found that the PDFG sequence is also conserved in peptidyl arginine deiminases (PADIs), and can assume two distinct conformations.^20^ Moreover, in certain kinase families the DFG motif is known to be part of a loop whose conformational states in many cases determine kinase activation. Taken together, these observations lead us to speculate that PDFG-like motifs may act as structural switches for multiple protein families.

## Results

### Long-timescale MD showed transient WPD loop states and candidate reaction coordinates for the WPD-loop transition

We performed multiple MD simulations starting from both the WPD-closed crystal structure 1SUG and the WPD-open crystal structure 2CM2. We observed the WPD loop stably transition from the closed state to open state four times in a total of 3.2 ms simulation time; we did not observe any transitions from the open state to the closed state. The fact that we observed the WPD loop transition in only one direction is consistent with the timescales derived from Whittier et al.’s NMR kinetic experiments, which suggested a timescale of ~45 ms for the transition from the open to the closed state.^6^ An example trajectory of the WPD loop transition is shown in Figure 1 and Movie S1; the WPD Loop transitions from the closed state to the open state at 78.31 μs. We note that our simulations oftentimes sampled a “transient open” state, wherein the canonical distance metric between the C215-Cα atom and D181-Cα atom^3^ increased for hundreds of nanoseconds but then reverted back to the closed state (yellow, Figure 1a). Similarly, even shorter-lived “transient closed” states were sampled (blue, Figure 1a), wherein the canonical distance metric between the C215-Cα atom and D181-Cα atom decreased below the 9.87 Å cutoff, indicating a closed state, but quickly reverted to the open state (Figure 1a). This suggested that the distance metric alone was not a sufficient reaction coordinate to capture the full transition of the WPD loop from one true, long-lived state to another.

**Figure 1.**
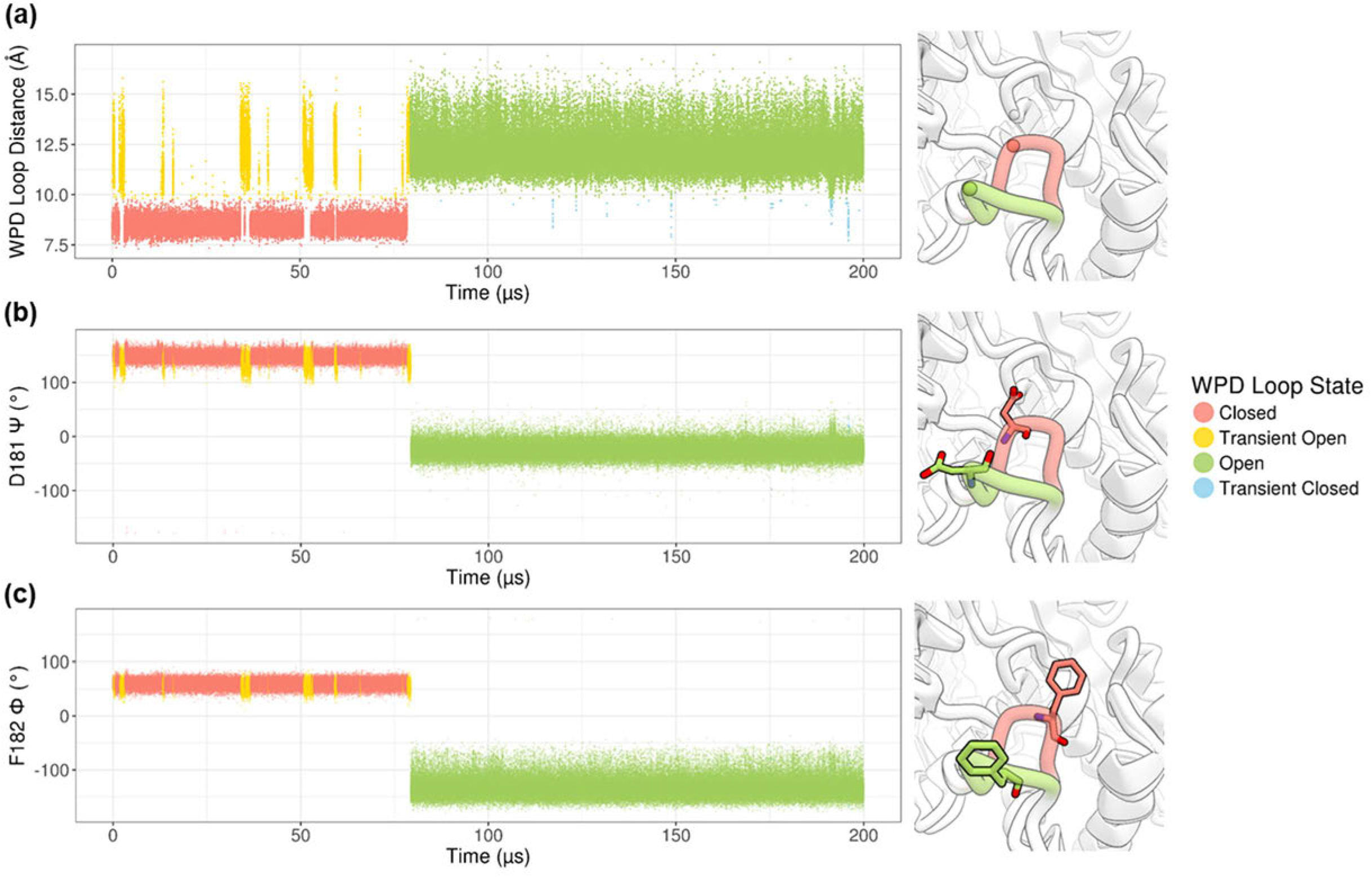
Long-timescale MD uncovered transient WPD loop states and candidate reaction coordinates. **(a)** The canonical D181-Cα to C215-Cα distance was observed as a reaction coordinate of the WPD loop transition from closed (red) to open (green). Sub-microsecond “transient open” states (yellow), and even shorter-lived “transient closed” states (blue), were revealed. **(b)** The D181-Ψ dihedral angle of the catalytic acid in the WPD loop was observed as a reaction coordinate of the WPD loop transition from closed (red) to open (green). **(c)** The F182-Φ dihedral angle of the phenylalanine in the WPD loop was captured as a reaction coordinate of the WPD loop transition from closed (red) to open (green).

Upon further interrogation of our trajectories, with correlational methods and dimension reduction analyses (Methods), we discovered many structural features correlated with the WPD loop transitioning from closed to open conformations (Figure S1). These included the Ψ dihedral in the backbone of the catalytic aspartate, D181, and the Φ dihedral in the backbone of the adjacent phenylalanine F182.^21^ We observed, as exemplified in Figure 1b and 1c, that these two quantitative measurements clearly demarcate the two long-lived WPD loop states.

### AWE simulations sampled the millisecond-timescale WPD loop transition

The above observations gave us the candidate degrees of freedom with which we could further investigate the atomic-level mechanism of the WPD loop catalytic transition. We collapsed our observations from unbiased, long-timescale MD simulations into two reaction coordinates, one of which measures the distance between the top of the loop and the active site, and the other of which measures a deformation within the loop. We refer to the former as “WPD Loop Distance (Å)” (distance between D181-Cα and C215-Cα atoms), and the latter as “WPD Loop RMSD (Å)” (RMSD of the backbone atoms of D181 and F182 to the reference 1SUG crystal structure). We verified that all wild-type PTP1B structures available in the Protein Data Bank (PDB) labeled either “closed” or “open” were well separated as two “states” in this two-dimensional space (Figure 2a). We also observed that the energy landscape, estimated from an aggregate 3.2 ms of MD simulations, also showed two energy wells that corresponded with clusters of PDB crystal structures along these two reaction coordinates.

**Figure 2.**
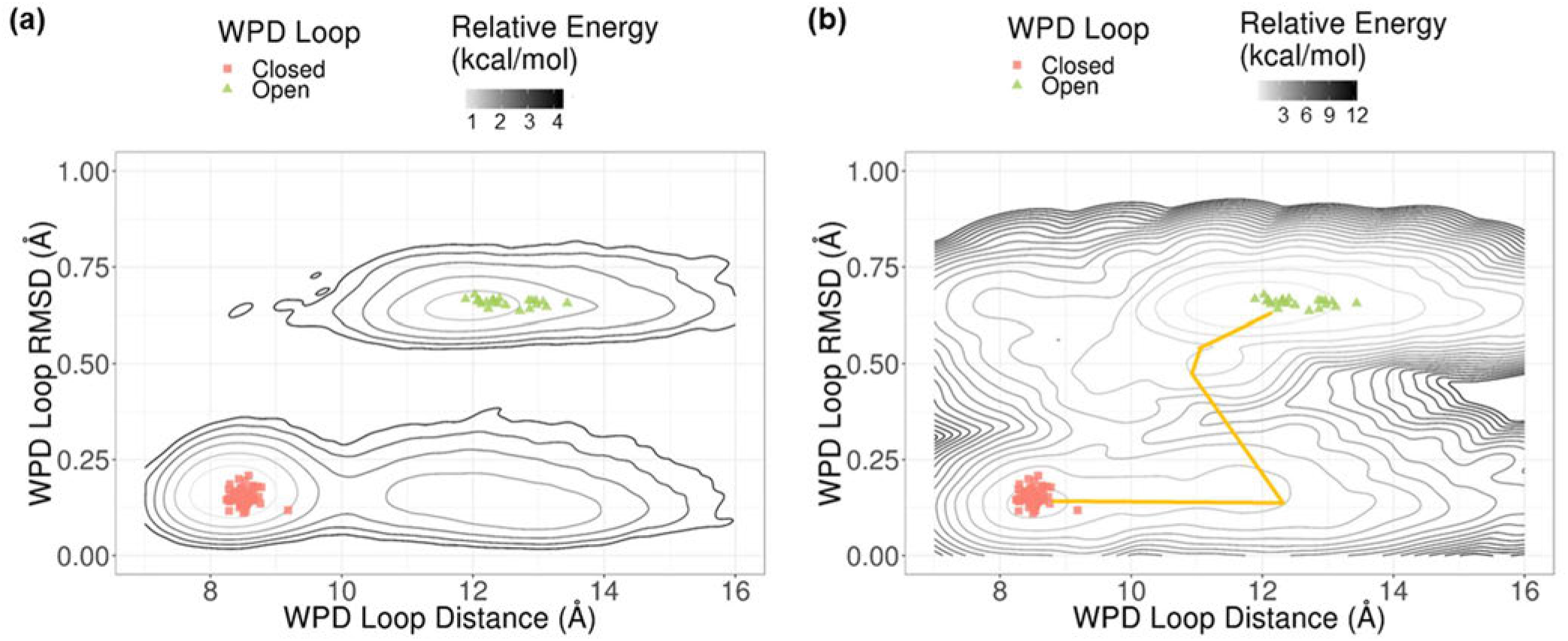
Kinetics and thermodynamics of the WPD loop transitions were accurately recapitulated in AWE simulations using two reaction coordinates obtained from inspection of long-timescale MD simulations and available crystal structures in the PDB. **(a)** The D181-Ψ and F181-Φ angles were condensed into the reaction coordinate “WPD Loop RMSD (Å).” The two reaction coordinates, “WPD Loop Distance (Å)” and “WPD Loop RMSD (Å),” separate the two states of the WPD Loop found in crystal structures of PTP1B in the PDB. Coarse energy estimates, using a total of 3.2 ms of simulation data, show two energy minima that correspond with the closed and open crystal structures from the PDB (each contour = 1 kcal mol^-1^). **(b)** Accelerated weighted ensemble (AWE) simulations yielded robust sampling of the millisecond WPD loop transition, with MFPT_closed-to-open_ = 4 ms, and MFPT_open-to-closed_ = 108 ms. Analysis of the AWE data using transition path theory^45,46^ results in the main flux line connecting five of the bins used during the AWE sampling (yellow line). This main flux line suggests a two-step mechanism of the WPD loop opening transition, similar to what we observed in unbiased MD simulation: D181-Cα –C215-Cα distance increases first, then the WPD loop dihedrals switch. The free energy difference between the “open” and “closed” states is −2.6 ± 0.1 kcal mol^-1^.

By sampling along these two reaction coordinates, “WPD Loop Distance (Å)” and “WPD Loop RMSD (Å),” our AWE simulation replicates converged and reproducibly recapitulated the millisecond-timescale WPD loop transitions from both closed to open and open to closed WPD loop states (Figure 2b; Methods). As shown in Figs. 2a and 2b, the locations and shapes of the basins in the unbiased MD and AWE simulations are consistent. The converged sampling in the AWE simulations gave us a wealth of kinetic and thermodynamic information. We computed the kinetics of the transitions: MFPT_closed-to-open_ = 4 ms and MFPT_open-to-closed_ = 108 ms, which are consistent with the experimentally determined^6^ kinetics of the WPD loop transition. The free energy estimate from these AWE simulations was ΔG_closed-to-open_ = −2.6 ± 0.1 kcal mol^-1^, indicating that the transition from closed to open states is spontaneous (Figure 2b), a finding that is again consistent with experimental data. The dominant flux of the WPD loop opening in the AWE simulations (Methods) follows a mechanism by which the WPD loop distance increases first, before motion largely in the WPD loop RMSD coordinate leads over an energy barrier to the closed state (Figure 2b).

### A parsimonious random forest model built on six backbone dihedrals of residues P180, D181, F182, and G183 captured the transition state ensemble of the WPD loop transition

In addition to recapitulating the kinetics and thermodynamics of the WPD loop transition, the replicate AWE simulations also yielded 1980 putative transition state structures (Figure 3a). We tested these putative transition state structures by computing their committor probabilities (P*_closed_*). We observed that the structures with estimated P*_closed_* = 0.5 ±0.1 did not occupy a distinct energy barrier in our estimated free energy landscape, suggesting that the reaction coordinates used are not entirely satisfactory for describing the WPD loop transition and in particular defining the transition states along it. We thus investigated whether we could obtain improved reaction coordinates using machine learning methods.

**Figure 3.**
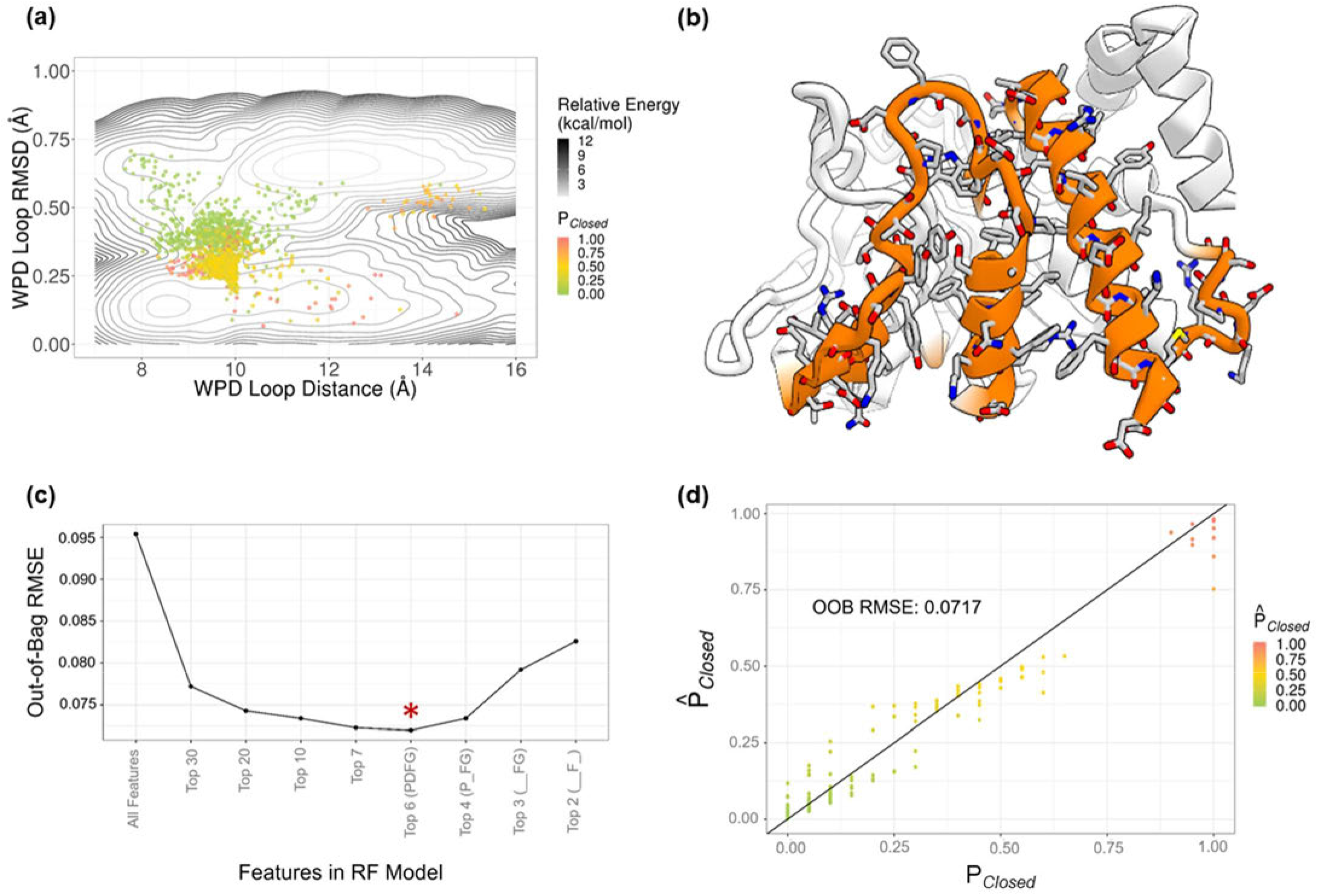
Feature pruning in random forest model selection yielded a parsimonious predictor of P*_Closed_* built on only six backbone dihedrals of residues P180, D181, F182, and G183 (PDFG model). **(a)** Putative transition state structures obtained from AWE simulations were tested on Anton for quantitative committor probability (P*_Closed_*). **(b)** Curated features reported in the literature (highlighted in orange, and heavy atoms explicitly shown in licorice) were used to build an initial random forest model to predict committor probability on putative TS structures. **(c)** Feature pruning shows that a model built on the top 6 features comprised of P180, D181, F182, G183 backbone dihedrals yields the lowest OOB RMSE (indicated by a red star). **(d)** The scatterplot of predicted committor probability 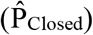 vs. calculated committor probability P_Closed_ shows the high predictive power of the PDFG model; the cross-validated model prediction error of the model was 0.0717 (3 s.f.)

We used feature pruning and model selection in the supervised machine learning method— random forest—to interrogate the contribution of a comprehensive set of structural features (highlighted in Figure 3b) to the dynamic mechanism of the WPD loop transition in PTP1B catalysis. By evaluating the performance of stepwise pruned models built on limited features, we determined that a model built on six dihedrals from the residues of P180, D181, F182, and G183 (starred, Figure 3c) was the most performant for accurately predicting P*_closed_*. We observed that in the six-dimensional space spanned by the selected six dihedrals from the residues of P180, D181, F182, and G183, structures with committor P*_closed_* = 0.5 ± 0.1 clustered separately from other putative transition-state structures (Figure S5), suggesting that these six backbone dihedrals provide a better reaction coordinate for describing the transition. The prediction accuracy of the final model, which we named the *PDFG model*, had an out-of-bag (OOB; see Methods) RMSE = 0.0717 (Figure 3d). (The OOB error is a bootstrap error estimate using the error of each training data point from the trees in the random forest that do not contain that point.) These results suggest that the feature space captured by these six PDFG backbone dihedrals is sufficient not only to correctly predict long-lived states of WPD-loop structures (P*_closed_* = 0, open and P*_closed_* = 1, closed), but also to accurately predict transition-state structures P*_closed_* = 0.5 ± 0.1) in out-of-sample test sets.

We further validated the PDFG model by showing that it can correctly predict the WPD loop states of sample structures drawn from the trajectory in Figure 1 (which was not included in the training of the model). The model predicted that all 30 frames drawn from the first 10 μs were closed 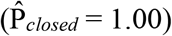, and all 30 frames drawn from 95–105 μs were open 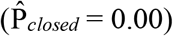. We also generated predictions for 1416667 frames between 78.48 μs and 79.33 μs, where we believe the true transition from the closed to open occurred in simulation. We note that the PDFG ensemble predictions did not shift to a mean above 0.5 as the WPD loop distance started to increase, but only after the switch in WPD loop RMSD (Figure S5). The model predicted that the four structures between 79310646 ps to 79310649 ps (emphasized in Figures 4b, S4, and S6) had committor probability 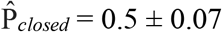. Of these four frames, all yielded non-trivial committor probabilities (P*_closed_*) between 0.6 and 0.9 in a follow-up committor analysis.

**Figure 4.**
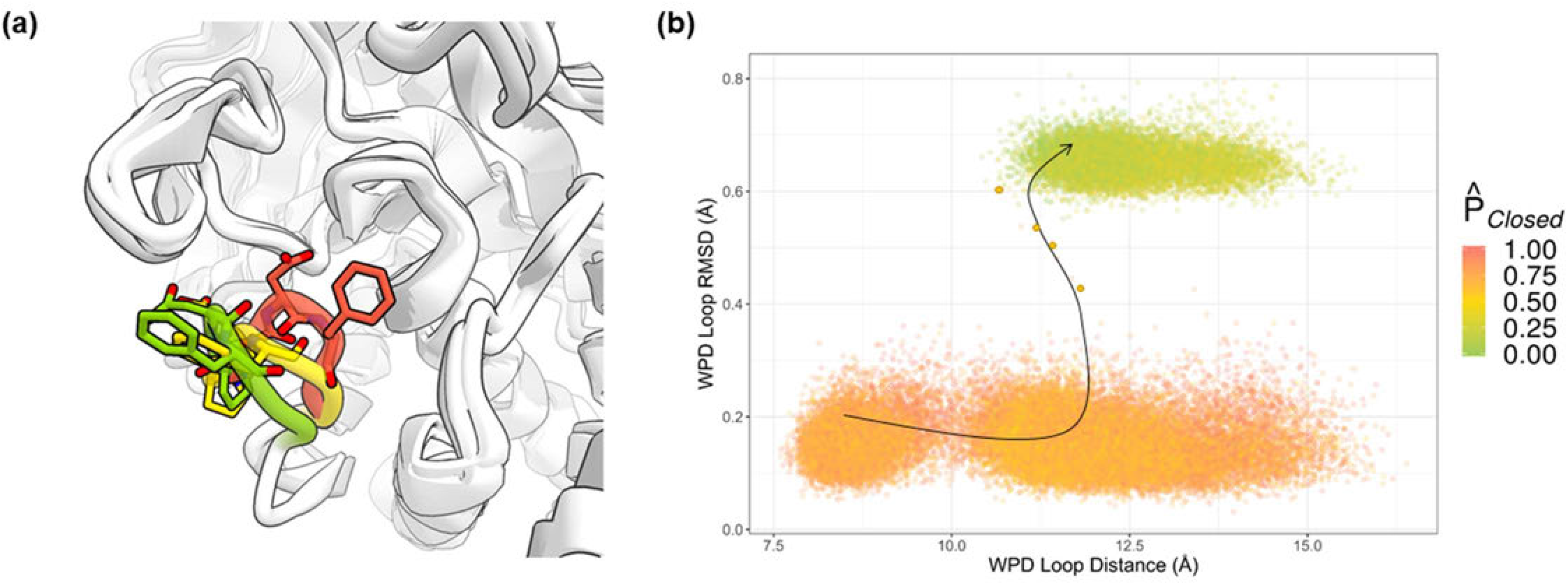
PDFG model predictions on an unbiased, long-timescale MD trajectory shows that the backbone dihedrals of the PDFG motif are sufficient to describe the WPD loop transition. **(a)** Representative structures of predicted closed (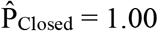, *t* = 0.23 μs, red), transition state (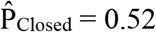, *t* = 78.31 μs, yellow), and open (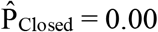, *t* = 100.78 μs, green) frames from the simulation. **(b)** The PDFG model correctly identifies the structures near the transition observed in the simulation as having intermediate committor probability (each dot in the plot represents a structure of the WPD loop obtained—at 6-ps intervals—from the simulation, and the four structures obtained during the transition are shown as four larger dots.) The black line qualitatively indicates the direction of the conformational transition observed in the MD simulation.

### PDFG and related sequences are conserved in different families of enzymes

In order to evaluate how widely applicable in the human genome these findings about the function of the PDFG motif might be, we performed an intra-family multiple sequence alignment (MSA) of all members of the PTP family, followed by an inspection of the available corresponding crystal structures in the PDB. Consistent with previous work, we found that the PD[F/H]G sequence is conserved in PTPs (Figure 5a).^2^ Structural alignment of available PTP crystal structures^22^ showed that there exist “open” and “closed” crystal structures of many of these phosphatases (Figure 5a), supporting the notion that the PD[F/H]G motif may also be an important structural switch for catalysis in other PTPs. A BLAST search on the PD[F/H]G motif also recovered matches to certain families of kinases, with the related “DFG” motif being fully conserved in these families.^23^ It is well known that the DFG motif adopts two conformational states in many kinase families—as, for example, in the ABL kinases (Figure 5c)—separating functionally active and inactive states, analogously to the role played by the PDFG motif in our model. Our PD[F/G]H BLAST search also revealed that the PDFG sequence is conserved in all but one protein arginine deiminase (PADI). Subsequent alignment and analysis of publicly available crystal structures of protein arginine deiminases also show two distinct conformational states of the loop at a Ca^2+^ binding site (Figure 5b). We note that although the PD[F/H]G BLAST search did return matches in other protein families, there was not the structural information corresponding to those matches that would be needed to draw further conclusions on the conformational significance of PD[F/H]G motifs in those families.

**Figure 5.**
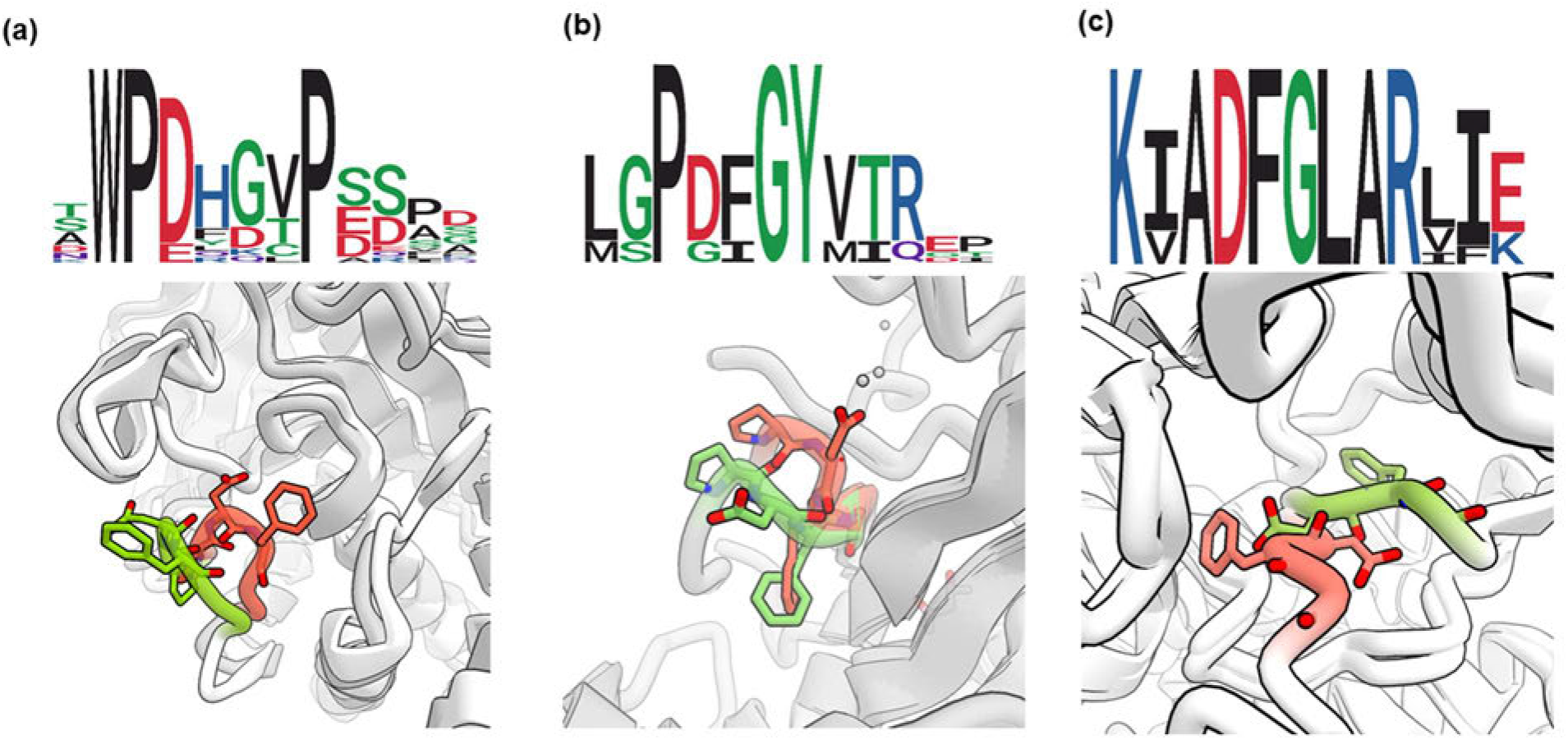
Multiple sequence alignment (MSA) and structural bioinformatic analysis suggests that PDFG-like motifs may be widely used as structural switches. **(a)** Sequence logo for PD[F/H]G motif in the PTP family of proteins and representative structures of crystallized PTP states: open (green, PDB: 2CM2) and closed (red, PDB: 1SUG). PDFG residues shown explicitly in licorice. **(b)** Sequence logo for the PDFG motif in PADIs and representative structures of crystallized PADI states: open (green, PDB: 4N20) and closed (red, PDB: 4N25). PDFG residues shown explicitly in licorice. **(c)** Sequence logo for the well-known DFG motif in the Src family of kinases and representative structures of crystallized ABL kinase states: DFG-in (green, PDB: 2F4J) and DFG-out (red, PDB: 1OPJ) DFG residues shown explicitly in licorice.

## Discussion

Although the atomic-level mechanism of the rate-limiting step of the transition has been unclear, numerous studies have made progress in identifying protein features associated with the conformational change of the WPD loop of PTP1B. Past studies have used approaches such as NMR, X-ray crystallography, MD simulations, and biochemistry, and made important observations of specific residues, interactions, and secondary structures associated with PTP1B’s catalytic function^24^ on distinct timescales.^10^ Others have used multi-temperature crystallography and chemical fragment probes^11^ to elucidate additional structural features that contribute to the functional regulation of PTP1B. Studies using protein NMR and crystallography have highlighted the similarities and differences in activity, structure, and dynamics between the respective WPD loops of PTP1B and its bacterial analogue, YopH, yielding additional features that could be involved in the mechanism of PTP1B’s catalytic control.^6,12^

In our study, we used simulations to discover and validate robust reaction coordinates of PTP1B’s catalytic cycle in several steps. First, unbiased, long-timescale simulations sampled several WPD loop transition events. Our sampling of these rare events allowed us to perform structural bioinformatic analyses to propose potential structural features as reaction coordinates, including D181Ψ and F182Φ. Subsequent AWE simulations then gave us statistically robust sampling of the WPD loop transition in both directions, and the analysis of this data helped yield a putative transition state ensemble that we later refined. We found that the PDFG backbone dihedrals are key to describing the rare but rapid transition from one long-lived state of the loop to another, and we conclude that that this motif thus acts as the structural arbiter of the WPD loop transition in PTP1B’s catalytic cycle.

This finding led us to perform a structural bioinformatic analysis of publicly available crystal structures: For PTPs, multiple kinase families, and PADIs, the PDFG-like sequence is conserved and confers a loop-like structure that exists in two distinct conformational states, analogously to the distinct “open” and “closed” states in PTP1B. This suggests that PDFG-like sequences act as structural switches for controlling catalysis in these protein families (and potentially others for which structural information is not yet available).

Knowledge of the function of this structural switch could have implications in the rational design of small-molecule drugs and in the design of enzymes: If the PDFG-like sequence indeed acts as a structural switch that controls enzymatic activity, one might in principle design small molecules that slow the activity of these enzymes by acting on the switch, in contrast with the typical approach of blocking the orthosteric binding site. (Past drug discovery programs have had success in modulating the enzyme when engaging the PDFG motif directly (in PADI2^27^) or indirectly (in PTP1B^8^), suggesting the potential promise of this approach.) It has been shown that just ~600 small tertiary structural motifs, similar in size to the PDFG motif, are sufficient to describe 50% of protein structures in the PDB at sub-Ångstrom resolution.^17,30^ Databases of such structural motifs have proved valuable in enzyme design,^31^ and we speculate that the functional role of the conserved PDFG-like motifs in separating long-lived protein states could potentially be exploited for the rational design of enzymes.

## Methods

### Long-timescale molecular dynamics

All PTP1B long-timescale simulations were performed on Anton,^32^ a special-purpose supercomputer for molecular dynamics simulations. These simulations were based on crystal structures 1SUG^33^ (“closed” state) and 2CM2^34^ (“open” state) from the PDB. Both constructs were truncated to include only residues 2–284. Histidines in the system were epsilon protonated. We solvated these structures in water with neutralizing NaCl counter ions at 150 mM in an 80 × 80 × 80 Å^3^ simulation box at 310 K. We used the Amber99SB*-ILDN^35^ protein force field (which builds on other modifications^36,37^ to Amber99^38^) with backbone dihedral and hydrogen bond restraints to stabilize the SBL (Tables S1 and S2), as described previously.^9^ For ions, the parameters of Aqvist^39^ (which are the default choice for Amber99SB*-ILDN) were used. The waters were parameterized with the TIP3P^40^ model. From the “open” state, we simulated 6 independently thermalized replicates for 200 μs each. From the “closed” state, we simulated 6 independently thermalized replicates for 200 μs each, and an additional 40 independently thermalized replicates for 20 μs each. The aggregate total simulation time was 3.2 ms.

### Accelerated weighted ensemble simulations

We performed all AWE simulations on Anton, using an in-house implementation of the AWE algorithm. Iterations of AWE consisted of two steps: First, unbiased MD was run for all currently active simulations (also called “walkers”) for a short time. In this work, all walkers were run for 100.8 ps, and their velocities were randomly initialized at every iteration. A stochastic thermostat was used, based on a Langevin dynamics integrator. The second step involved a two-color resampling: Walkers that started from an “open” state had an “open” color property, and walkers that started from a “closed” state had a “closed” color property. When walkers of a given color reached the opposite state, they switched colors. Resampling was performed separately on each walker-color population using an algorithm similar to those described in Costagueo et al.^13^ and Abdul-Wahid et al.^41^ Walkers were split when they carried too much probability weight compared to a target weight, and they were merged when they carried too little weight.

Initially, an AWE simulation was started from the “open” state. We picked two reaction coordinates that were discretized as follows: the “WPD Loop Distance (Å)” (distance between D181-Cα and C215-Cα atoms) used four bins, and the “WPD Loop Distance (Å)” (distance between D181-Cα and C215-Cα atoms) used fourteen bins (Figure S2). Each occupied bin was assigned 20 walkers, and each walker ran for 100 ps. The first simulation ran for 78 iterations— 40 μs aggregate simulation time—until transitions in both directions were collected: 751 open to closed, and 3394 closed to open. The fluxes in both directions achieved steady state (roughly equal). A Markov state model (MSM) was formed based on the observed transition probabilities during the first simulation, and it yielded equilibrium populations of 0.97 for the “open” state and 0.02 for the “closed” state, corresponding to a free energy difference of −k_B_T*log(P*_open_*/P*_closed_*), or −2.3 kcal mol^-1^. To obtain more detailed kinetics and snapshots of the mechanism, further simulations were run from this endpoint, starting with the MSM-reweighted bins. The rates were computed as

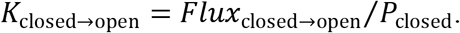

Here, the flux is time-averaged, and computed using only walkers with the “closed” color to avoid overcounting; similarly, the probability includes only walkers with the “closed” color. Likewise,

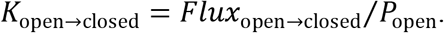

Error bars for the free energy and the rates were obtained by running 3 replicates of 30 iterations each—~6 μs per replicate—for an aggregate simulation time of 18 μs.

### Committor probability analysis (P_closed_)

We computed committor probability (P*_closed_*) estimates for each putative transition state structure extracted from the AWE simulations. Starting from each structure, we performed 20 randomly thermalized simulations until each simulation either reached the closed state (WPD Loop Distance < 8.689 Å, WPD Loop RMSD < 0.269 Å) or open state (WPD Loop Distance > 10.075 Å, WPD Loop RMSD > 0.590 Å). We calculated *P_closed_* for each putative transition state structure as the ratio between the number of trajectories that committed to the closed state and the total number of runs (20).

### Random forest and model selection

A set of 361 structural features associated with the PTP1B catalytic mechanism was curated from the PTP1B literature. We then computed these features for each putative transition state from the AWE simulations. A 10-fold cross-validated, bootstrap-aggregated (bagged) random forest model was then trained on these features to predict P*_closed_* as the response variable 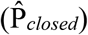.

Feature importance scores were averaged across 10 folds (Figure S3) and assigned a rank *k* in increasing order, where *k* = 1 denotes the most important feature, and *k* = 361 the least important feature. A descendent strategy,^42^ with incrementally smaller numbers of the top features, were used to build new models (Figure 3c). We converged on a parsimonious, yet performant, model by evaluating each pruned model using the OOB error estimate.

### BLAST and multiple sequence alignment

We submitted the query “PD[F/H]G” in Standard Protein BLAST (blastp) with the BLOSUM62 algorithm and default parameters for short sequences. In the resulting list, each protein hit was mapped to UniProt. Entries with shared PFAM annotations were grouped together. The full amino acid sequences for each protein family group with more than one protein was then submitted to T-Coffee Multiple Sequence Alignment^43^ for full sequence alignments and quantitative probabilities on specific amino acid conservation. The resulting alignment and scores were then used to generate sequence logos with the “ggseqlogo”^44^ R package.

## Acknowledgments

The authors thank Yibing Shan for drawing our attention to the DFG motif of PTP1B, Fabrizio Giordanetto, Sahar Shahamatdar, Hunter Nisonoff, and Ellen Zhong for helpful discussions, and Berkman Frank and Eric Martens for editorial assistance.

## Notes

### Competing Interest Statement

The authors have declared no competing interest.

