## Supplemental Information for "A conserved local structural motif controls the kinetics of PTP1B catalysis"

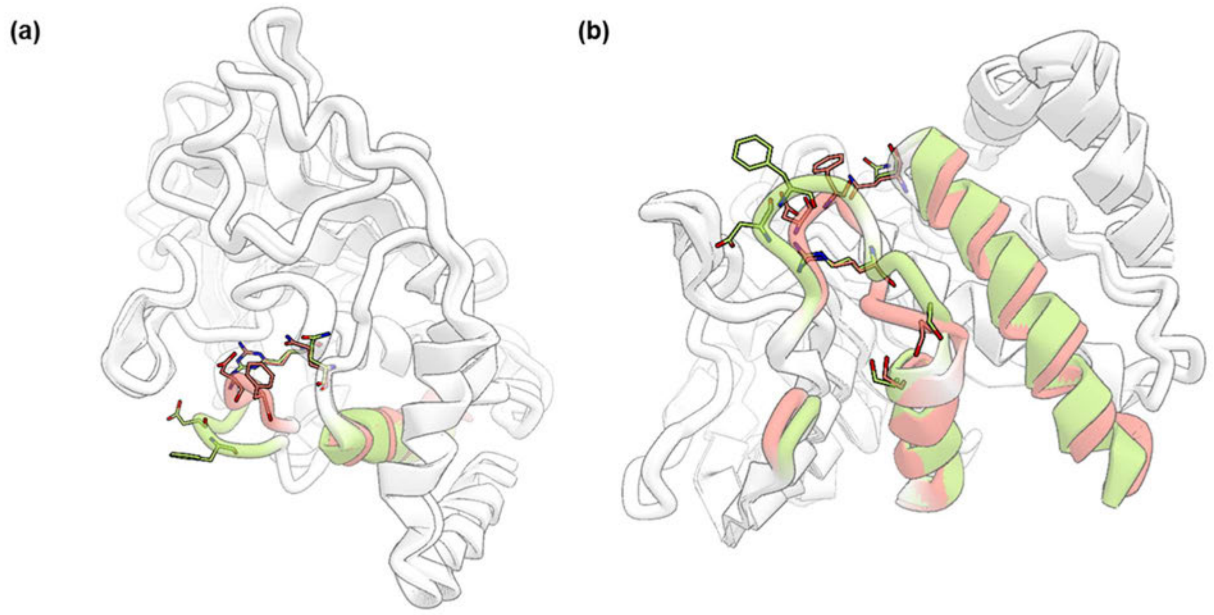


**Figure S1. Specific structural features were found to be correlated with the WPD Loop transition in unbiased, long-timescale MD simulations. (a)** Residues and secondary structures correlated with WPD loop closed (red) and open (green) states (top view). **(b)** Residues and secondary structures correlated with WPD loop closed (red) and open (green) states (front view).

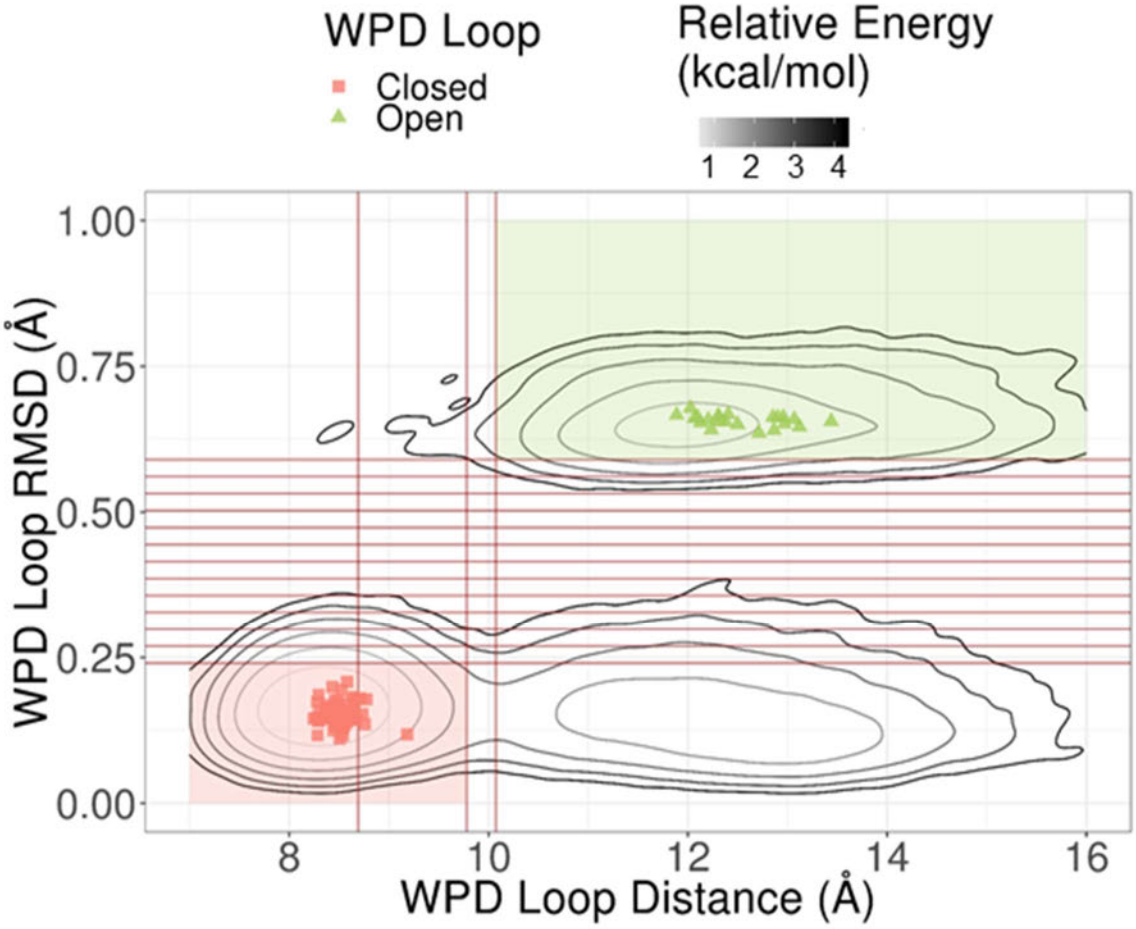


**Figure S2. Analyses of long-timescale simulations and available PDB crystal structures of PTP1B informed the binning and state definitions in designing AWE simulations.**

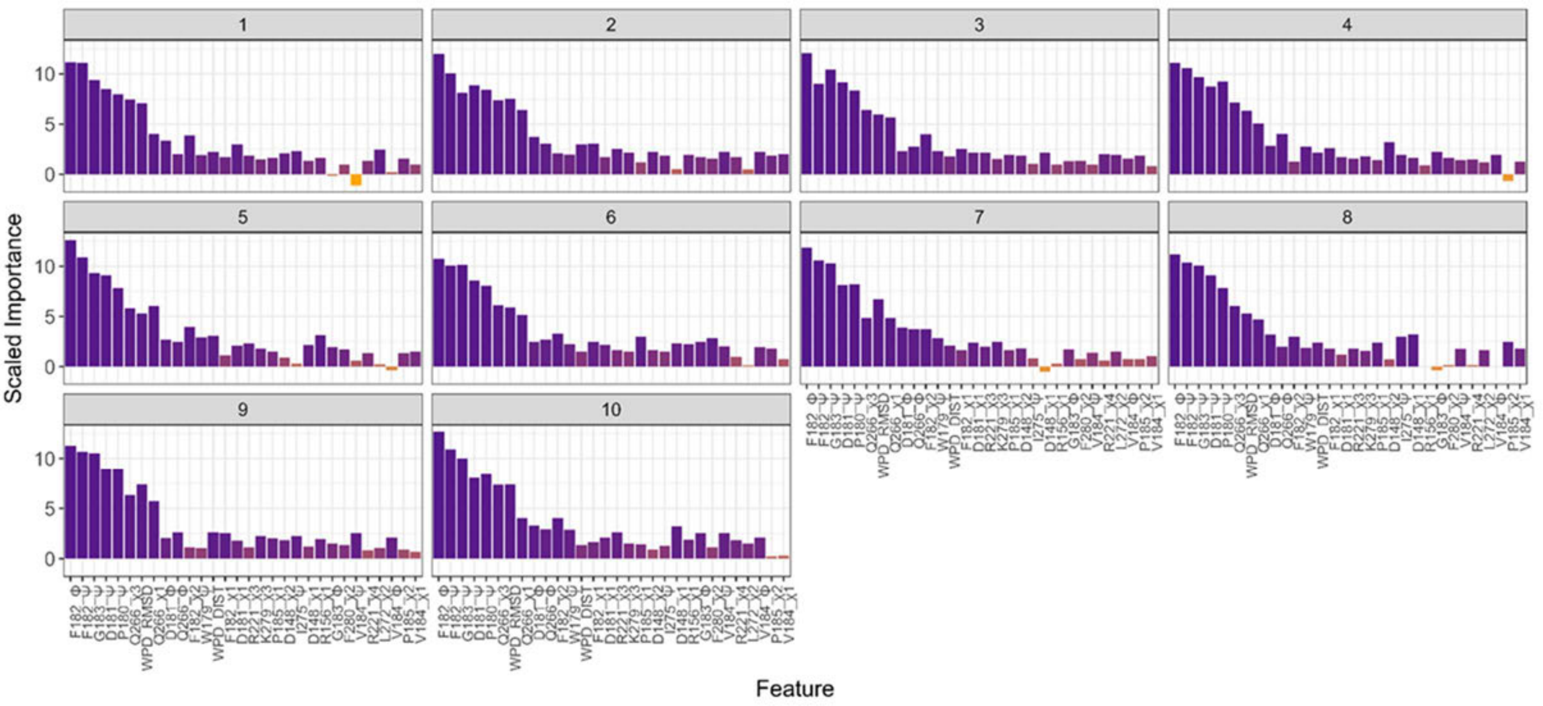


**Figure S3. Feature Importance Ranking in 10-fold cross-validation of the global Random Forest model is robust across all folds for top 30 features.**

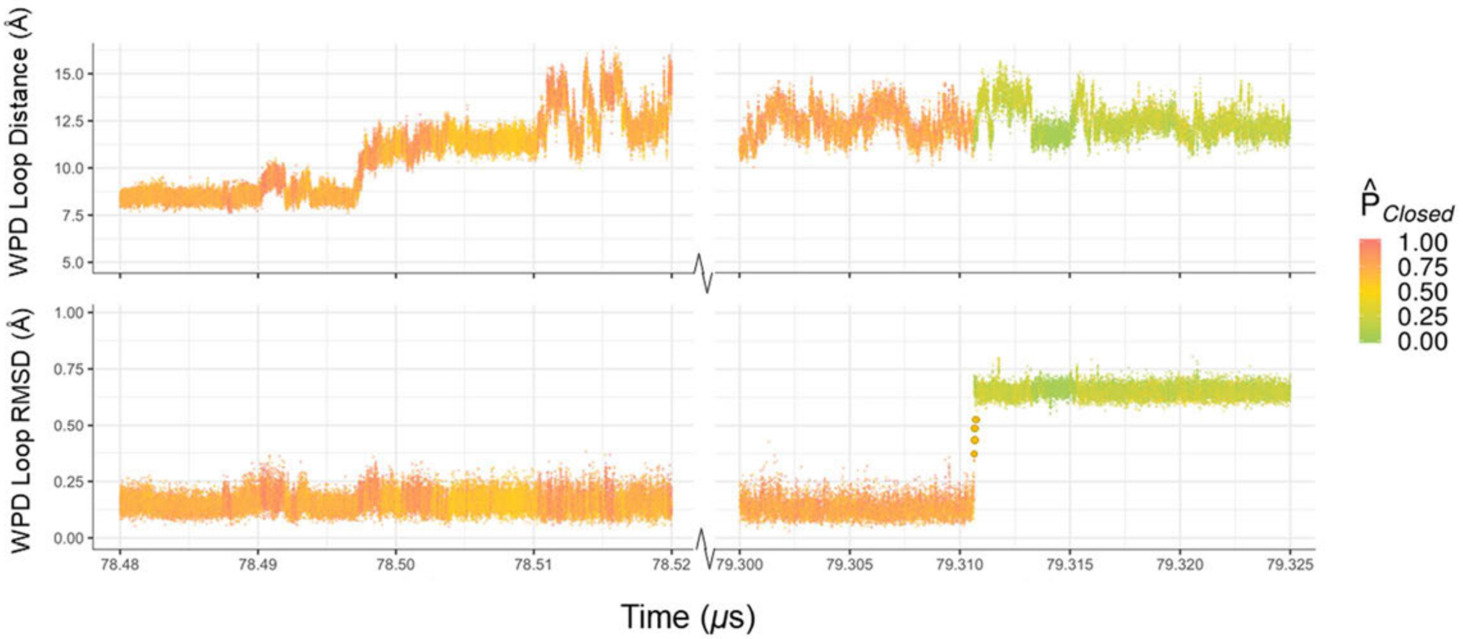


**Figure S4. The PDFG model predictions on an unbiased, long-timescale MD trajectory capture an exact switch between PTP1B end-states.** The structures along the transition path are emphasized by larger yellow dots.

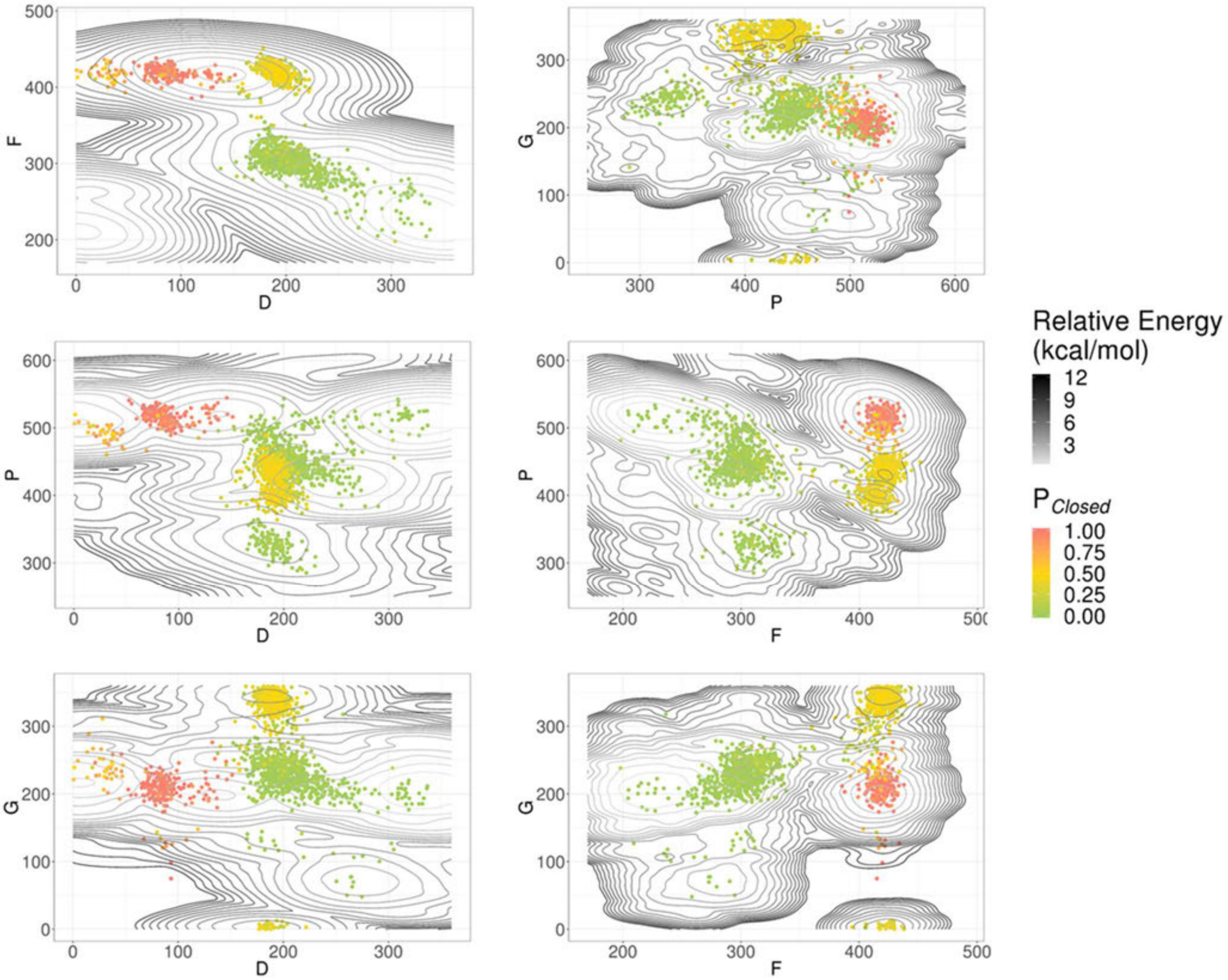


**Figure S5. Putative TS structures with $P_{Closed}$ ~0.5 occupy distinct energy wells in PDFG dihedral space.**

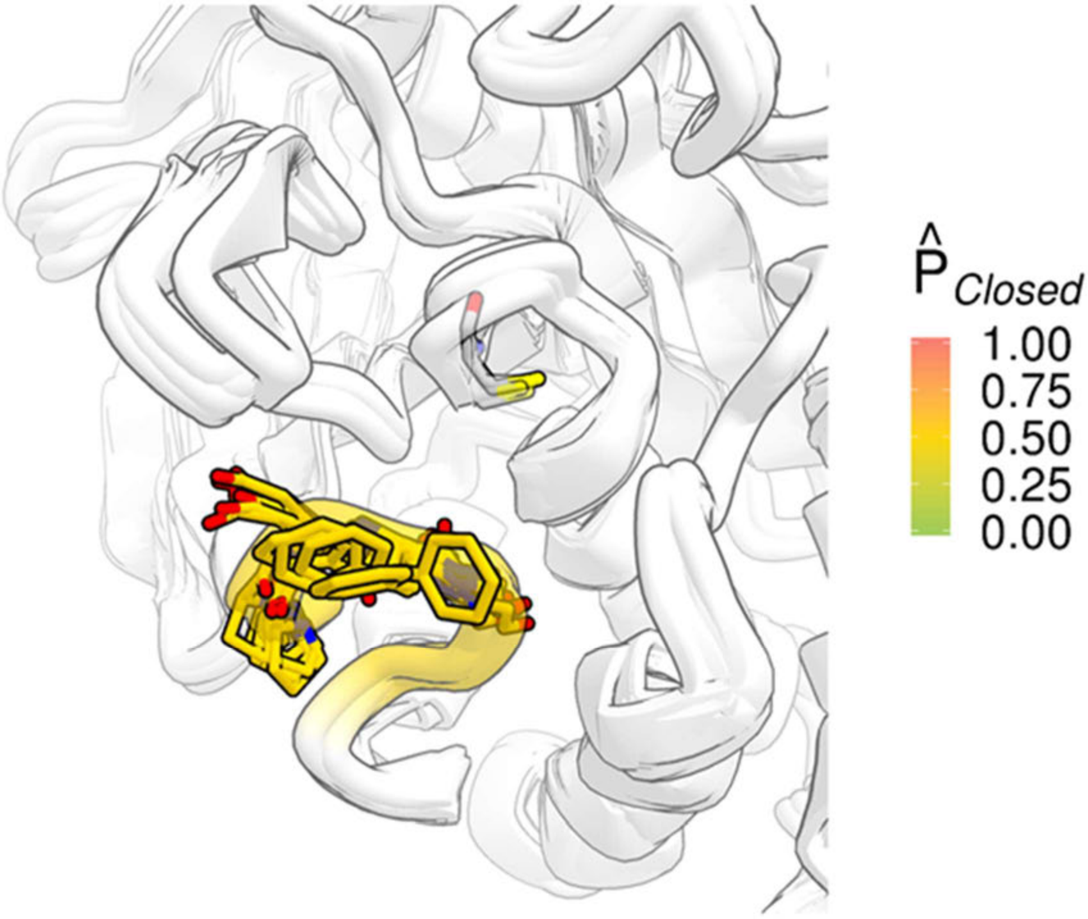


**Figure S6. Four superimposed, highlighted snapshots of the structures along the transition path from the apo MD simulation shown in Figure S4.** These structures have predicted committor probability $0.45 < \hat{P}_{Closed} < 0.55$ using the committor-prediction model with 6 dihedral angles from the residues P180, D181, F182, and G183. These four residues are shown explicitly in licorice without hydrogens and are colored based on the PDFG-model predictions ($\hat{P}_{closed}$).

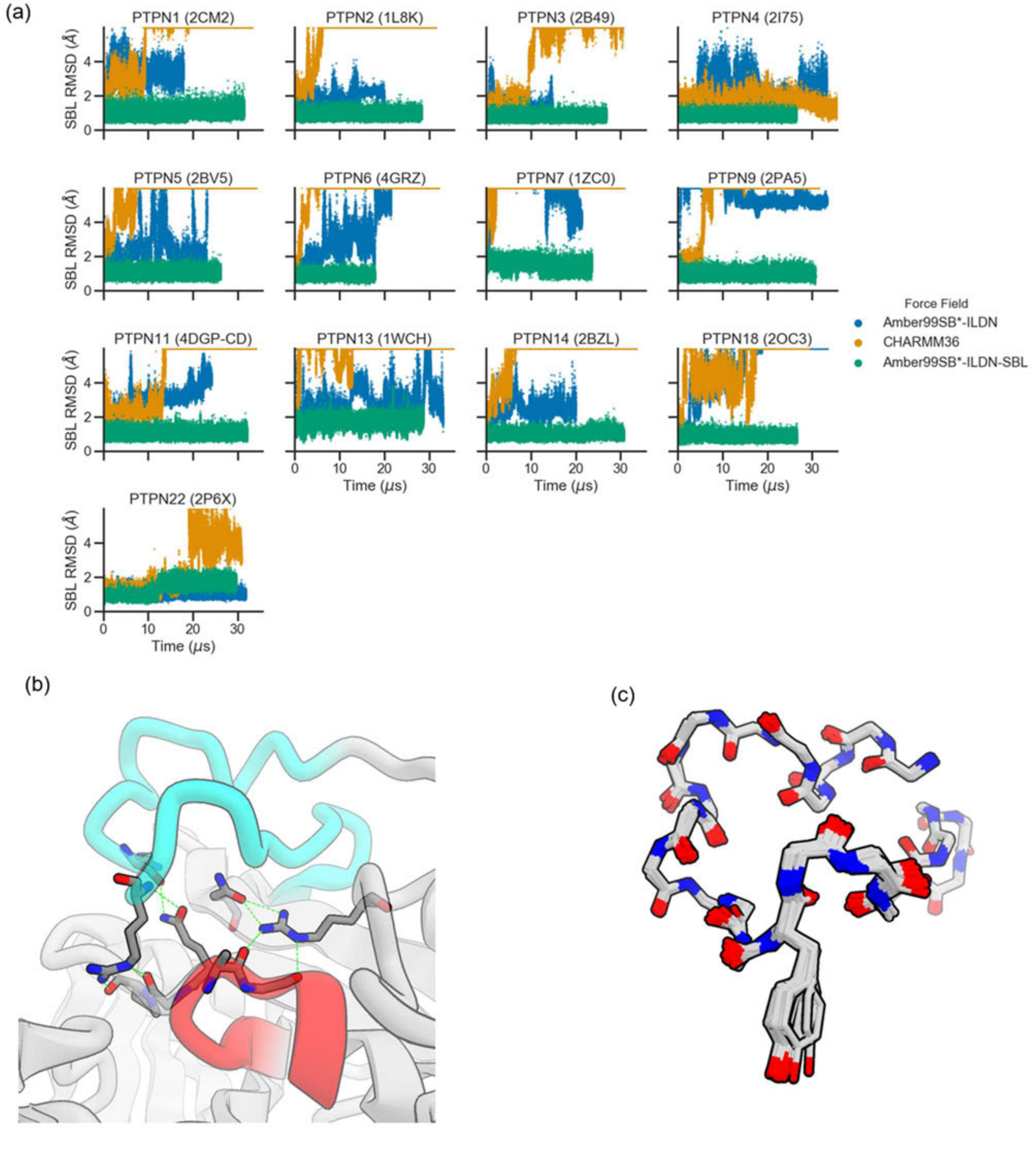


**Figure S7. The substrate-binding loop (SBL) of non-receptor PTPs was unstable on microsecond timescales in common force fields.** (a) MD simulations of non-receptor PTPs were run in Amber99SB*-ILDN, CHARMM36, and Amber99SB*-ILDN with restraints stabilizing the SBL. Cα-RMSD of the SBL after alignment to a conserved selection from the core of the PTP fold was high (>5 Å) for simulations with Amber99SB*-ILDN and CHARMM36, but low for simulations with SBL restraints. (b) Hydrogen bonds included in the SBL restraints are shown with green dashed lines. (c) Overlay of the backbone atoms of the SBL for all deposited PTP1B structures with wild-type sequence.

| PTPN | ATOM1 | ATOM2 |
|---|---|---|
| 1 | ARG45 NE | GLY86 O |
| 1 | ARG45 NH2 | PRO87 O |
| 1 | GLN85 NE2 | ARG43 O |
| 1 | GLN85 OE1 | ARG45 N |
| 1 | ARG257 NH2 | ALA217 O |
| 1 | ARG257 NE | GLY218 O |
| 1 | ARG257 NH2 | ASN68 OD1 |
| 1 | ARG257 NH1 | ASN68 OD1 |
| 2 | ARG47 NE | GLY88 O |
| 2 | ARG47 NH2 | PRO 89 O |
| 2 | GLN87 NE2 | ARG47 O |
| 2 | GLN87 OE1 | ARG47 N |
| 2 | ARG255 NH2 | ALA218 O |
| 2 | ARG255 NE | GLY219 O |
| 2 | ARG255 NH2 | ASN70 OD1 |
| 2 | ARG255 NH1 | ASN70 OD1 |
| 3 | ARG675 NE | GLY719 O |
| 3 | ARG675 NH2 | PRO 720 O |
| 3 | GLN718 NE2 | LYS673 O |
| 3 | GLN718 OE1 | ARG675 N |
| 3 | ARG881 NH2 | ALA844 O |
| 3 | ARG881 NE | GLY845 O |
| 3 | ARG881 NH2 | ASN697 OD1 |
| 3 | ARG881 NH1 | ASN697 OD1 |
| 4 | ARG684 NE | GLY728 O |
| 4 | ARG684 NH2 | PRO729 O |
| 4 | GLN727 NE2 | LYS682 O |
| 4 | GLN727 OE1 | ARG684 N |
| 4 | ARG891 NH2 | ALA854 O |
| 4 | ARG891 NE | GLY855 O |
| 4 | ARG891 NH2 | ASN706 OD1 |
| 4 | ARG891 NH1 | ASN706 OD1 |
| 5 | ARG303 NE | GLY350 O |
| 5 | ARG303 NH2 | PRO351 O |
| 5 | GLN349 NE2 | LYS301 O |
| 5 | GLN349 OE1 | ARG303 N |
| 5 | ARG511 NH2 | ALA473 O |

| | | |
|---|---|---|
| 5 | ARG511 NE | GLY474 O |
| 5 | ARG511 NH2 | ASN331 OD1 |
| 5 | ARG511 NH1 | ASN331 OD1 |
| 6 | ARG275 NE | GLY326 O |
| 6 | ARG275 NH2 | CYS327 O |
| 6 | GLN325 NE2 | LYS273 O |
| 6 | GLN325 OE1 | ARG275 N |
| 6 | ARG495 NH2 | ALA455 O |
| 6 | ARG495 NE | GLY456 O |
| 6 | ARG495 NH2 | ASN303 OD1 |
| 6 | ARG495 NH1 | ASN303 OD1 |
| 7 | ARG103 NE | GLY149 O |
| 7 | ARG103 NH2 | PRO150 O |
| 7 | GLN148 NE2 | LYS101 O |
| 7 | GLN148 OE1 | ARG103 N |
| 7 | ARG309 NH2 | ALA272 O |
| 7 | ARG309 NE | GLY273 O |
| 7 | ARG309 NH2 | ASN130 OD1 |
| 7 | ARG309 NH1 | ASN130 OD1 |
| 9 | ARG332 NE | GLY377 O |
| 9 | ARG332 NH2 | PRO378 O |
| 9 | GLN376 NE2 | LYS330 O |
| 9 | GLN376 OE1 | ARG332 N |
| 9 | ARG554 NH2 | ALA517 O |
| 9 | ARG554 NE | GLY518 O |
| 9 | ARG554 NH2 | ASN359 OD1 |
| 9 | ARG554 NH1 | ASN359 OD1 |
| 11 | ARG278 NE | GLY332 O |
| 11 | ARG278 NH2 | CYS333 O |
| 11 | GLN331 NE2 | LYS276 O |
| 11 | GLN331 OE1 | ARG278 N |
| 11 | ARG501 NH2 | ALA461 O |
| 11 | ARG501 NE | GLY462 O |
| 11 | ARG501 NH2 | ASN306 OD1 |
| 11 | ARG501 NH1 | ASN406 OD1 |
| 13 | ARG2242 NE | GLY2284 O |
| 13 | ARG2242 NH2 | PRO2285 O |
| 13 | GLN2283 NE2 | LYS2240 O |

| | | |
|---|---|---|
| 13 | GLN2284 OE1 | ARG2242 N |
| 13 | ARG2447 NH2 | ALA2410 O |
| 13 | ARG2447 NE | GLY2411 O |
| 13 | ARG2447 NH2 | ASN2264 OD1 |
| 13 | ARG2447 NH1 | ASN2264 OD1 |
| 14 | ARG938 NE | GLY984 O |
| 14 | ARG938 NH2 | PRO985 O |
| 14 | GLN983 NE2 | ARG936 O |
| 14 | GLN983 OE1 | ARG938 N |
| 14 | ARG1160 NH2 | ALA1123 O |
| 14 | ARG1160 NE | GLY1124 O |
| 14 | ARG1160 NH2 | ASN964 OD1 |
| 14 | ARG1160 NH1 | ASN964 OD1 |
| 18 | ARG61 NE | GLY106 O |
| 18 | ARG61 NH2 | PRO107 O |
| 18 | GLN105 NE2 | LYS59 O |
| 18 | GLN105 OE1 | ARG61 N |
| 18 | ARG271 NH2 | ALA231 O |
| 18 | ARG271 NE | GLY232 O |
| 18 | ARG271 NH2 | ASN88 OD1 |
| 18 | ARG271 NH1 | ASN88 OD1 |
| 22 | ARG59 NE | GLY104 O |
| 22 | ARG59 NH2 | PRO105 O |
| 22 | GLN103 NE2 | LYS57 O |
| 22 | GLN103 OE1 | ARG59 N |
| 22 | ARG269 NH2 | ALA229 O |
| 22 | ARG269 NE | GLY230 O |
| 22 | ARG269 NH2 | ASN86 OD1 |
| 22 | ARG269 NH1 | ASN86 OD1 |

**Table S1.** Distance restraints of the SBL that enforce the hydrogen bond network. The first column shows the gene, and the second and third columns show the pair of atoms that form the hydrogen bond.

| PTPN | RESIDUES |
|---|---|
| 1 | 32-56 |
| 2 | 34-58 |
| 3 | 662-686 |
| 4 | 671-695 |
| 5 | 314-338 |
| 6 | 262-286 |
| 7 | 111-135 |
| 8 | 319-343 |
| 11 | 265-289 |
| 13 | 2229-2253 |
| 14 | 925-949 |
| 18 | 48-72 |
| 22 | 46-70 |

**Table S2.** Backbone dihedral restraints of the SBL. The first column shows the gene, and the second column the range of residues that is restrained.

**Movie S1. Long-timescale molecular dynamics (MD) shows transient WPD loop states and candidate reaction coordinates.** In the movie, the WPD loop is colored by the discrete "states" of the loop. The coloring of the loop corresponds to the color scheme in Figure 1 (closed, red; transient open, yellow; open, green; transient closed, blue).
